# Identification of Host Biomarkers of EBV Latency IIb and Latency III

**DOI:** 10.1101/616359

**Authors:** Joshua E. Messinger, Joanne Dai, Lyla J. Stanland, Alexander M. Price, Micah A. Luftig

**Affiliations:** Department of Molecular Genetics and Microbiology, Duke Center for Virology, Duke University School of Medicine Durham, NC, USA

## Abstract

Deciphering the molecular pathogenesis of virally induced cancers is challenging due, in part, to the heterogeneity of both viral and host gene expression. Epstein-Barr Virus (EBV) is a ubiquitous herpesvirus prevalent in B-cell lymphomas of the immune suppressed. EBV infection of primary human B cells leads to their immortalization into lymphoblastoid cell lines (LCLs) serving as a model of these lymphomas. In previous studies, our lab has described a temporal model for immortalization with an initial phase characterized by expression of the Epstein-Barr Nuclear Antigens (EBNAs), high c-Myc activity, and hyper-proliferation in the absence of the Latent Membrane Proteins (LMPs), called latency IIb. This is followed by the long-term outgrowth of LCLs expressing the EBNAs along with the LMPs, particularly the NFκB-activating LMP1, defining latency III. LCLs, however, express a broad distribution of LMP1 such that a subset of these cells expresses LMP1 at levels seen in latency IIb, making it difficult to distinguish these two latency states. In this study, we performed mRNA-Seq on early EBV-infected latency IIb cells and latency III LCLs sorted by NFκB activity. We found that latency IIb transcriptomes clustered independently from latency III independent of NFκB. We identified and validated mRNAs defining these latency states. Indeed, we were able to distinguish latency IIb cells from LCLs expressing low levels of LMP1 using multiplex RNA-FISH targeting EBV EBNA2, LMP1, and human CCR7. This study defines latency IIb as a *bona fide* latency state independent from latency III and identifies biomarkers for understanding EBV-associated tumor heterogeneity.

**IMPORTANCE:** EBV is a ubiquitous pathogen with >95% of adults harboring a life-long latent infection in memory B cells. In immunocompromised individuals, latent EBV infection can result in lymphoma. The established expression profile of these lymphomas is latency III, which includes expression of all latency genes. However, single cell analysis of EBV latent gene expression in these lymphomas suggests heterogeneity where most cells express the transcription factor, EBNA2, and only a fraction express the membrane protein LMP1. Our work describes an early phase after infection where the EBNAs are expressed without LMP1, called latency IIb. However, LMP1 levels within latency III vary widely making these states hard to discriminate. This may have important implications for therapeutic responses. It is crucial to distinguish these states to understand the molecular pathogenesis of these lymphomas. Ultimately, better tools to understand the heterogeneity of these cancers will support more efficacious therapies in the future.

## INTRODUCTION

Epstein-Barr Virus (EBV) is a large double-stranded DNA γ-herpesvirus that establishes life-long latent infection in resting memory B cells. Despite robust immune control in the vast majority of infected individuals, immune-compromised patients are at high risk for EBV-driven B-cell lymphomas. A model for these lymphomas is EBV infection and immortalization of primary human B cells *in vitro* into lymphoblastoid cell lines (LCLs). Immortalized LCLs express all eight EBV latency proteins, consistent with latency III gene expression, including the EBV nuclear antigen (EBNA) transcription factors and the latent membrane proteins (LMPs), which are constitutively-active receptor mimics (1-3).

However, EBV-infected B cells initially undergo a period of hyper-proliferation characterized by expression of the EBNAs in the near absence of the LMPs, which is called latency IIb (4, 5). Early after infection, EBNA2 stimulates cellular proliferation by inducing the host transcription factor c-Myc through coordination of its upstream enhancer and chromatin looping (6). During this period, the cells are dependent upon MCL-1 and BCL-2 for survival in the absence of NFκB signaling (5, 7). Elevated levels of c-Myc early after infection antagonize LMP1 mRNA and protein expression (8). Low LMP1 levels early after infection may function to evade CD8+ T-cell recognition as LMP1-mediated NFκB activity promotes MHC-I expression and peptide presentation (9, 10). By two to three weeks post-infection, c-Myc levels wane, hyper-proliferation is attenuated, and full expression of the LMPs, particularly LMP1-mediated NFκB activity, is observed (11). These cells rely on NFκB signaling for survival (12) and display a distinct mitochondrial anti-apoptotic phenotype with up-regulation of BFL-1 (5, 7).

While latency III is characterized by full expression of the latent membrane proteins, it has long been observed that LMP1 levels vary widely within an LCL population. Flow cytometric analysis for LMP1 within bulk LCL populations shows an ∼100 fold range of LMP1 protein levels at single-cell resolution (13). This variable expression appears to be important for LCL homeostasis as significantly elevated or depleted LMP1 levels results in reduced proliferation and cells sorted for high or low levels of LMP1 return to their full distribution within 18 hours of sorting (10, 13). Therefore, LMP1 expression within an LCL population fluctuates widely on a single-cell level and this wide distribution is important for LCL survival.

EBV is associated with several different lymphomas including Hodgkin’s Lymphoma, Burkitt Lymphoma and Post-Transplant Lymphoproliferative Disease (PTLD). However, the viral latency gene expression in EBV-associated diseases are typically very heterogeneous. To understand the latency gene expression pattern in these diseases, immunohistochemical staining is employed to analyze the expression of LMP1 and EBNA2 in patient biopsies. Staining patient biopsies has demonstrated heterogeneity at the single cell level where many cells may be EBNA2-positive but negative for LMP1 (14, 15). These cells are often quite common as recent studies in a mouse model of EBV and Kaposi Sarcoma Herpes Virus (KSHV) co-infection also identified a high frequency of EBNA2+/LMP1-cells (16). Due to the wide distribution of LMP1 expression within a latency III LCL population, this technique does not allow for distinguishing LMP1 low latency III LCLs from LMP1 low latency IIb cells.

We have previously demonstrated that latency IIb and latency III cells have unique survival requirements and apoptotic regulation (7, 17, 18). However, these studies analyzed bulk LCL populations and did not address single-cell level differences. In this study we address such single-cell heterogeneity by fluorescence-activated cell sorting (FACS) sorting latency III LCLs based on surface ICAM-1 as a proxy for LMP1-mediated NFκB activity. Using this sorting strategy, we explored whether latency IIb cells are unique from a subset of NFκB-low LCLs that express reduced levels of LMP1. We also identified host transcriptomic markers of these latency states that are expressed in a latency stage dependent manner, but independent of LMP1 expression levels. Taken together, this work characterizes latency IIb as a unique B-cell latency state of EBV infection and identifies biomarkers that enables discrimination between latency IIb and latency III.

## RESULTS

### ICAM-1 is a proxy for LMP1-mediated NFκB target gene activation

We have previously demonstrated that LMP1 levels are significantly lower early after EBV infection of primary human B cells than in immortalized LCLs (5). However, it is also known that LMP1 levels vary widely within an LCL population (10, 13). Due to the wide distribution of LMP1 expression in an LCL population, we first asked how these levels compare to early-infected proliferating latency IIb cells. To assay this at the protein level, we conducted FACS analysis on early proliferating latency IIb cells (day 7 post-infection) and donor matched LCLs (>35 days post-infection) for ICAM-1 as a proxy for LMP1 mediated NFκB activity (19). As described in **Fig. 1A**, a subset of LCLs display similarly low levels of NFκB activity to latency IIb cells, corroborating our previous data. Therefore, we concluded that these cells likely express comparable levels of LMP1 mRNA. However, we wondered whether these LMP1^lo^ LCLs were unique from latency IIb cells, or cells that were “stuck” in latency IIb. To test this, we sorted latency IIb cells to purity as well as the bottom, middle and upper 15% of ICAM-1 expressing cells within donor matched LCL populations (**Fig. 1A**). We used RT-qPCR analysis to validate that ICAM-1 levels were similar between latency IIb cells and ICAM-1^lo^ LCLs and that ICAM-1 mRNA abundance increased with increasing ICAM-1 MFI (**Fig. 1B**). Importantly, LMP1 mRNA abundance followed the same pattern as ICAM-1 (**Fig. 1C**). In fact, there was a direct linear correlation between ICAM-1 and LMP1 mRNA abundance, thereby validating our sorting strategy (**Fig. 1D**).

**Figure 1:**
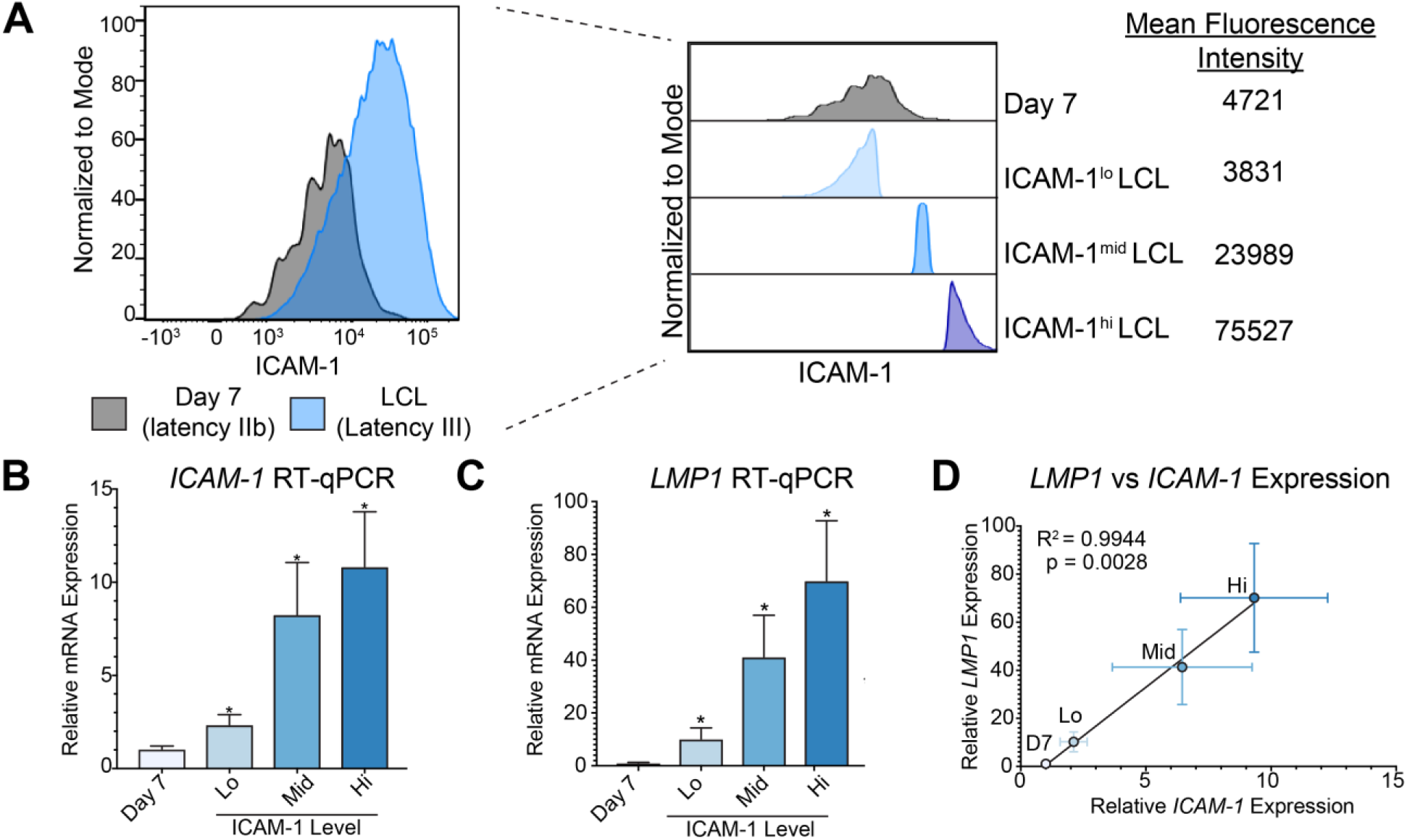
Using ICAM-1 as a proxy for LMP1 expression and LMP1-mediated NFκB signaling. A) FACS analysis for ICAM-1 as a proxy for LMP1-mediated NFκB activity of day 7 dpi EBV infected latency IIb proliferating B-cells and donor matched LCLs. Offset plot shows sorting groups of latency IIb as well as the bottom, middle and upper 15% of ICAM-1 expressing LCLs with corresponding MFI B) RT-qPCR for ICAM-1 in sorted groups. Each bar represents the average of six independent matched donors. C) RT-qPCR for LMP1 in sorted groups. Each bar represents the average of six independent matched donors. D) Correlation between LMP1 RT-qPCR expression and ICAM-1 RT-qPCR expression. * p<0.05, ** p<0.01, *** p<0.001 by student’s pairwise t-test. All error bars denote SEM.

### Generation and validation of RNA-Seq libraries from EBV early-infected and LCL populations sorted on ICAM-1 level

We next generated RNA-sequencing libraries to asses global gene expression differences between these four populations: B cells 7 days post EBV infection (latency IIb) and LCLs with low, middle, or high levels of ICAM-1 (**Fig. 1A**). We first sought to validate our RNA-Seq libraries by querying the global gene expression differences between ICAM-1^lo^ and ICAM-1^hi^ LCLs. Consistent with our initial sorting and qPCR experiments, we found by GSEA analysis a significant enrichment in NFκB targets in ICAM-1^hi^ relative to ICAM-1^lo^ LCLs (**Fig. 2A**). Indeed, RNA-Seq coverage maps indicate that two well-described NFκB targets, TRAF1 and A20, are expressed at higher levels in ICAM-1^hi^ relative to ICAM-1^lo^ LCLs (**Fig. 2B**). RT-qPCR analysis confirmed the RNA-seq results (**Fig. 2C**) and these data suggest internal validation of both our sorting approach and our RNA-Seq pipeline.

**Figure 2:**
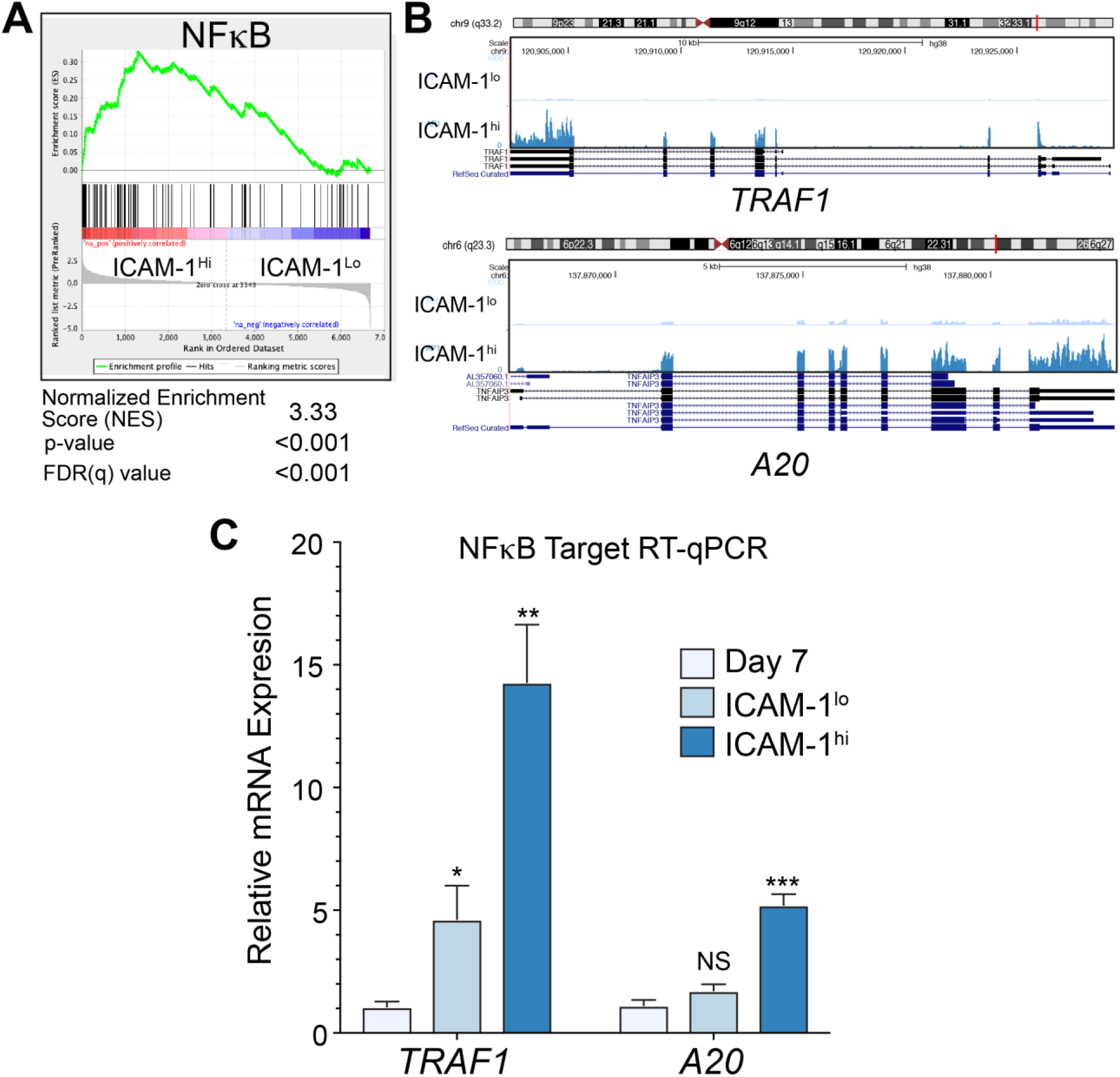
LCL populations are homogeneous despite wide a wide LMP1/NFκB distribution. A) Motif Pre-Ranked GSEA analysis between ICAM-1^lo^ and ICAM-1^hi^ expressing LCLs. B) RNA-seq coverage map at known NFκB targets TRAF1 and A20 for ICAM-1^lo^ and ICAM-1^hi^ LCLs C) RT-qPCR for TRAF1 and A20 in donor matched day 7 dpi EBV infected proliferating latency IIb cells and ICAM-1^lo^ or ICAM-1^hi^ LCLs. Each bar represents the average of six independent matched donors. * p<0.05, ** p<0.01, *** p<0.001 by student’s pairwise t-test. All error bars denote SEM.

### Host genes that distinguish EBV latency IIb cells from ICAM-1^lo^ LCLs are associated with DNA replication

Our major goal in this study is to define the host genes that distinguish early-infected latency IIb cells from ICAM-1^lo^ LCLs. Therefore, we performed a direct comparison of the genes differentially expressed between these two populations and identified 192 genes that were up-regulated in the transition from early-infected latency IIb cells to ICAM-1^lo^ LCLs relative and 216 genes down-regulated from latency IIb to ICAM-1^lo^ LCLs (**Fig. 3A** and **Supp. Table S1 and S2**). We performed GSEA for transcription factor motifs upstream of the differentially expressed genes and found that E2F family transcription factors as well as MYC/MAX transcription factors were significantly enriched in latency IIb compared to ICAM-1^lo^ LCLs (**Fig. 3B**). GSEA also identified several gene ontology groups associated with DNA replication and mitotic division as the hallmark of latency IIb as compared to ICAM-1^lo^ LCLs (**Fig. 3C** shows representative plot). To validate these findings, we interrogated expression of the genes associated with DNA replication by qRT-PCR. We found that MCM10, RFC2, RAD51, and PCNA were consistently, but not significantly, upregulated in latency IIb as compared to ICAM-1^lo^ LCLs (**Fig. 3D**). Furthermore, this difference was not observed between latency IIb and ICAM-1^mid^ or ICAM-1^hi^ LCLs. Given our inability to identify host markers that distinguish latency IIb from latency III from within this gene ontology group, we next sought to query the RNA-Seq data more broadly to identify such markers.

**Figure 3:**
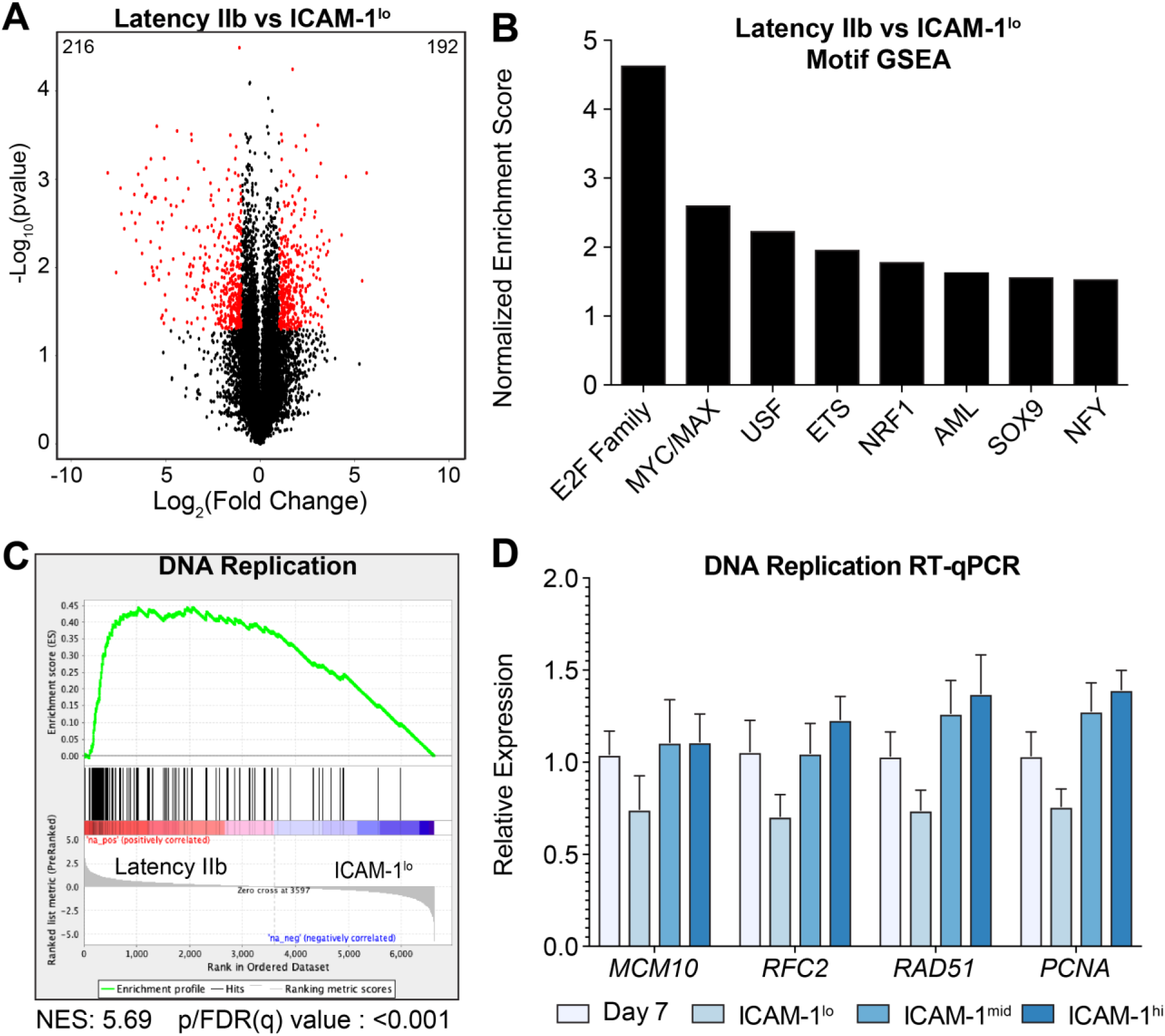
Latency IIb is defined by hyper-proliferation and enhanced DNA replication. A) Volcano plot of gene expression between latency IIb and ICAM-1^lo^ LCLs. Significantly regulated genes are indicated with red dots and have p<0.05 and a log_2_(Fold Change) of greater than 1 or less than negative 1. The number in the top left indicates the number of significantly regulated genes in latency IIb and the number in the top right denotes the number of significantly regulated genes for ICAM-1^lo^ B) Pre-ranked motif GSEA analysis between latency IIb and ICAM-1^lo^ LCLs C) Motif Pre-ranked GSEA between latency IIb and ICAM-1^lo^ LCLs for DNA replication D) RT-qPCR between latency IIb and ICAM-1^lo^ LCLs for DNA-replication genes. Each bar represents the average of six independent donors. Numbers on top of bars indicate the number of times this motif was listed by GSEA. * p<0.05, ** p<0.01, *** p<0.001 by student’s pairwise t-test. All error bars denote SEM.

### EBV early-infected latency IIb cells are transcriptomically distinct from latency III LCLs irrespective of ICAM-1/LMP1 levels

We next sought to assess whether the transcriptome of latency IIb cells varied significantly from the ICAM-1 sorted LCL populations. We first generated a Pearson coefficient similarity matrix comparing the expression profiles of all 16 samples. Day 7 latency IIb transcriptomic profiles clustered together independently of donor and the ICAM-1/LMP1 expression of the donor (**Fig. 4A**). The remaining clusters were comprised of the ICAM-1^lo^, ICAM-1^mid^, and ICAM-1^hi^ groups with each donor clustering independently from the others. These results were further substantiated by unsupervised hierarchical clustering of the samples where we found that day 7 latency IIb cells clustered independently of latency III LCLs (**Fig. 4B**). Therefore, gene expression differences between latency IIb and latency III cells are greater than those between donors and also between latency IIb and any ICAM-1 sorted population.

**Figure 4:**
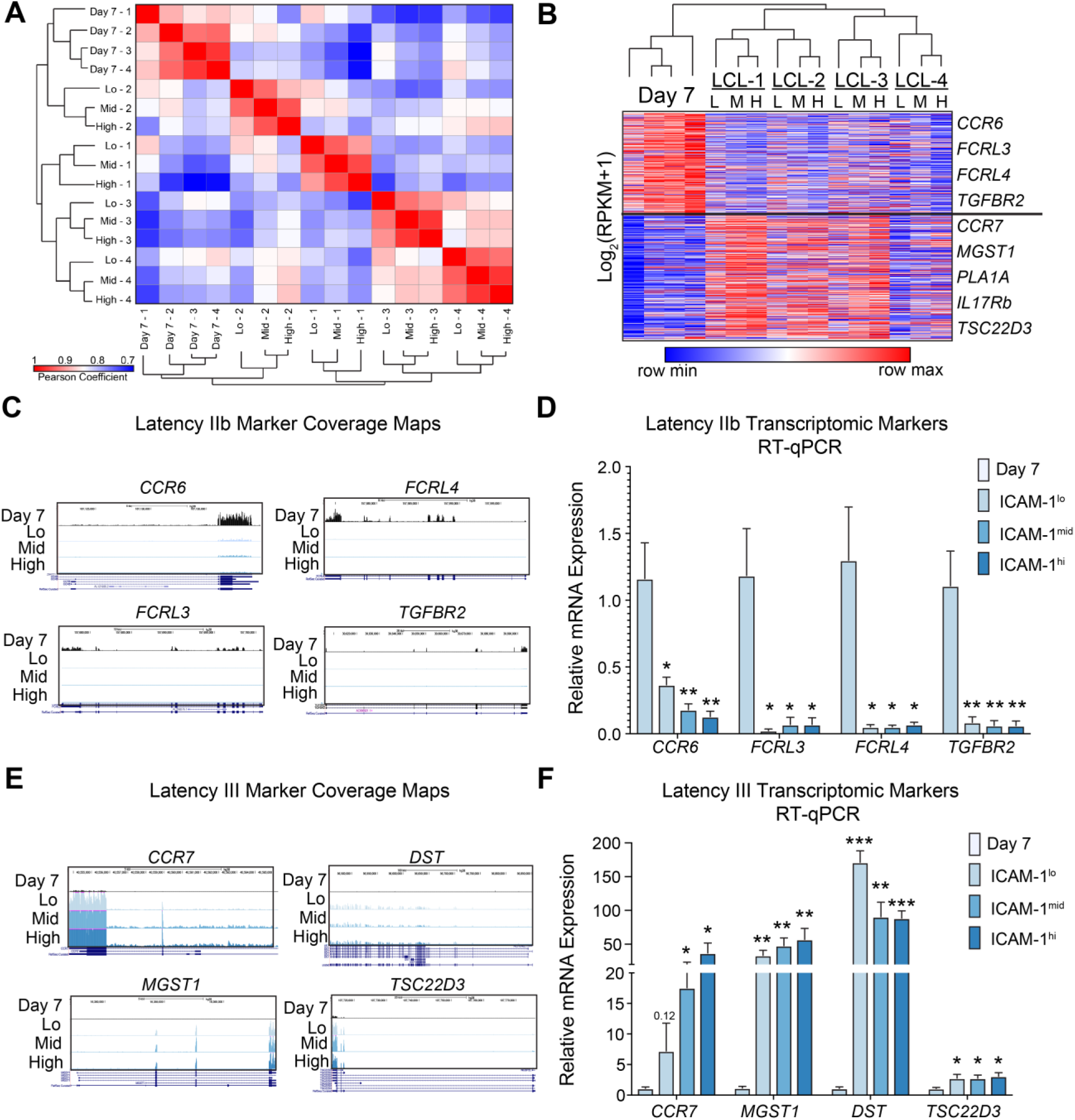
Latency IIb clusters uniquely from latency III independent of donor and both states contain transcriptionally unique host markers. A) Pearson coefficient similarity matrix and hierarchical clustering between all samples used for RNA-sequencing B) Gene expression heatmap with hierarchical and k-means clustering of gene expression data from RNA-sequencing C) RNA-sequencing coverage maps for latency IIb specific identified genes D) RT-qPCR validation of latency IIb specific genes E) RNA-sequencing coverage maps for latency III specific identified genes E) RT-qPCR validation of latency III specific genes. Each bar represents the average of six independent matched donors. * p<0.05, ** p<0.01, *** p<0.001 by student’s pairwise t-test. All error bars denote SEM.

K-means clustering of the gene expression data generated profiles uniquely associated with latency IIb and latency III. Within these profiles, we identified significantly differentially expressed genes based on a greater than 2-fold change and p<0.05 in at least two of the three comparison (day 7 versus ICAM-1^lo^, ICAM-1^mid^, or ICAM-1^hi^ LCLs). This analysis yielded 181 latency IIb-specific and 282 latency III-specific genes (**Supp. Table S3**). We chose four genes from each group with binary-like expression behavior to validate their specificity to latency IIb or latency III. Host biomarkers of latency IIb were *CCR6, FCRL3, FCRL4*, and *TGFBR2*. RNA-Seq coverage maps illustrate and qRT-PCR experiments validated IIb specificity of these genes (**Fig. 4C-D**). Latency III biomarkers were *CCR7, MGST1, DST*, and *TSC22D3* and these displayed similar binary-like gene expression behavior (**Fig. 4E-F**).

### Analysis of CCR6 and CCR7 surface expression as markers of latency IIb and III, respectively

CCR6 and CCR7 displayed the strongest expression differences between latency IIb and latency III by RNA-Seq (**Fig. 4C** and **4E**). As these proteins are both surface-expressed, we chose to investigate their utility as protein biomarkers to distinguish latency IIb from latency III. Flow cytometry of CCR6 indicated a strong down-regulation of surface expression comparing day 7 post infection to LCLs irrespective of ICAM-1 level corroborating our RNA-Seq and qRT-PCR data (**Fig. 5A**). While the mean fluorescence intensity (MFI) for CCR6 decreased significantly from day 7 to LCL (**Fig. 5B**), the difference in the percent of positive cells only dropped by half between day and ICAM-1lo LCLs (**Fig. 5C**). Similarly, while surface expression of CCR7 increased from day 7 to LCL (**Fig. 5D-E**), the percent positive cells only increased about two-fold (**Fig. 5F**). These data suggest that CCR6 and CCR7 protein levels will not suffice to distinguish between latency IIb and latency III expressing cells.

**Figure 5:**
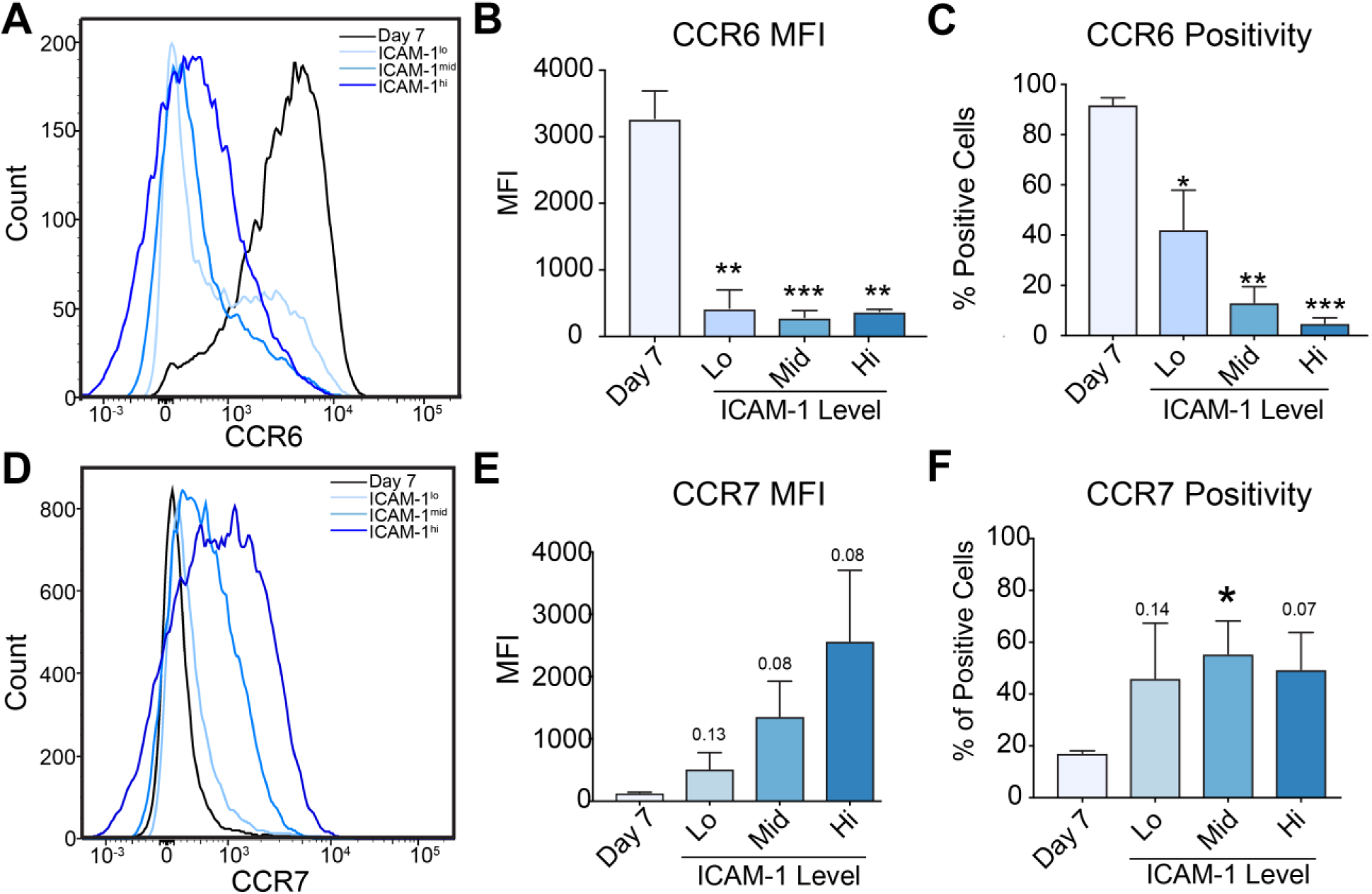
Protein validation of transcriptional markers reveals non-binary latency specificity. A) FACS analysis for CCR6 in day 7 latency IIb proliferating B cells and latency III LCLs stratified by ICAM-1 expression B) Quantification of mean fluorescence intensity of CCR6 expression from A C) Percentage of CCR6 positive cells from A D) FACS analysis for CCR7 in day 7 latency IIb proliferating B cells and latency III LCLs stratified by ICAM-1 expression E) Quantification of mean fluorescence intensity of CCR7 expression from D F) Percentage of CCR6 positive cells from D. Each bar represents the average of 3 independent donors * p<0.05, ** p<0.01, *** p<0.001 by student’s pairwise t-test. All error bars denote SEM.

### Multiplex RNA-FISH can distinguish latency IIb from latency III

Given the challenges of protein-based biomarker validation, we sought to use multiplex RNA fluorescence *in situ* hybridization (RNA-FISH) to leverage our RNA-based biomarker discovery approach. As detailed in **Fig. 4E**, the mRNA expression of *CCR7* strongly correlated with latency III independent of ICAM-1/LMP1 levels. CCR7 was also the most highly abundant latency III-specific mRNA. Therefore, we designed probes to detect CCR7 mRNA along with EBNA2 and LMP1 mRNAs. Our hypothesis would predict that latency IIb cells (EBNA2^+^/LMP1^-^) would be CCR7-negative and that latency III cells would be CCR7-positive irrespective of LMP1 level. We tested this hypothesis in sorted, proliferating day 7 (latency IIb) infected cells and LCLs with resting B cells and the EBV-negative B-lymphoma cell line, BJAB, as negative controls (**Fig. 6A).** EBNA2 expression was robust in EBV-infected day 7 cells and LCLs as compared to resting B cells and BJABs (**Fig. 6A-B)**. LMP1 expression was low in day 7 and higher, but heterogeneous, in LCLs, as expected (**Fig. 6A-C**). CCR7 levels were low in EBNA2+ day 7 cells and significantly higher in LCLs independent of LMP1 level (**Fig. 6D-E)**. Indeed, the full heterogeneity of LMP1 expression levels in latency III is visualized in **Fig. 6E** where all cells are EBNA2-positive and CCR7-positive. Thus, CCR7 mRNA is a reliable host transcriptomic marker of EBV latency III.

**Figure 6:**
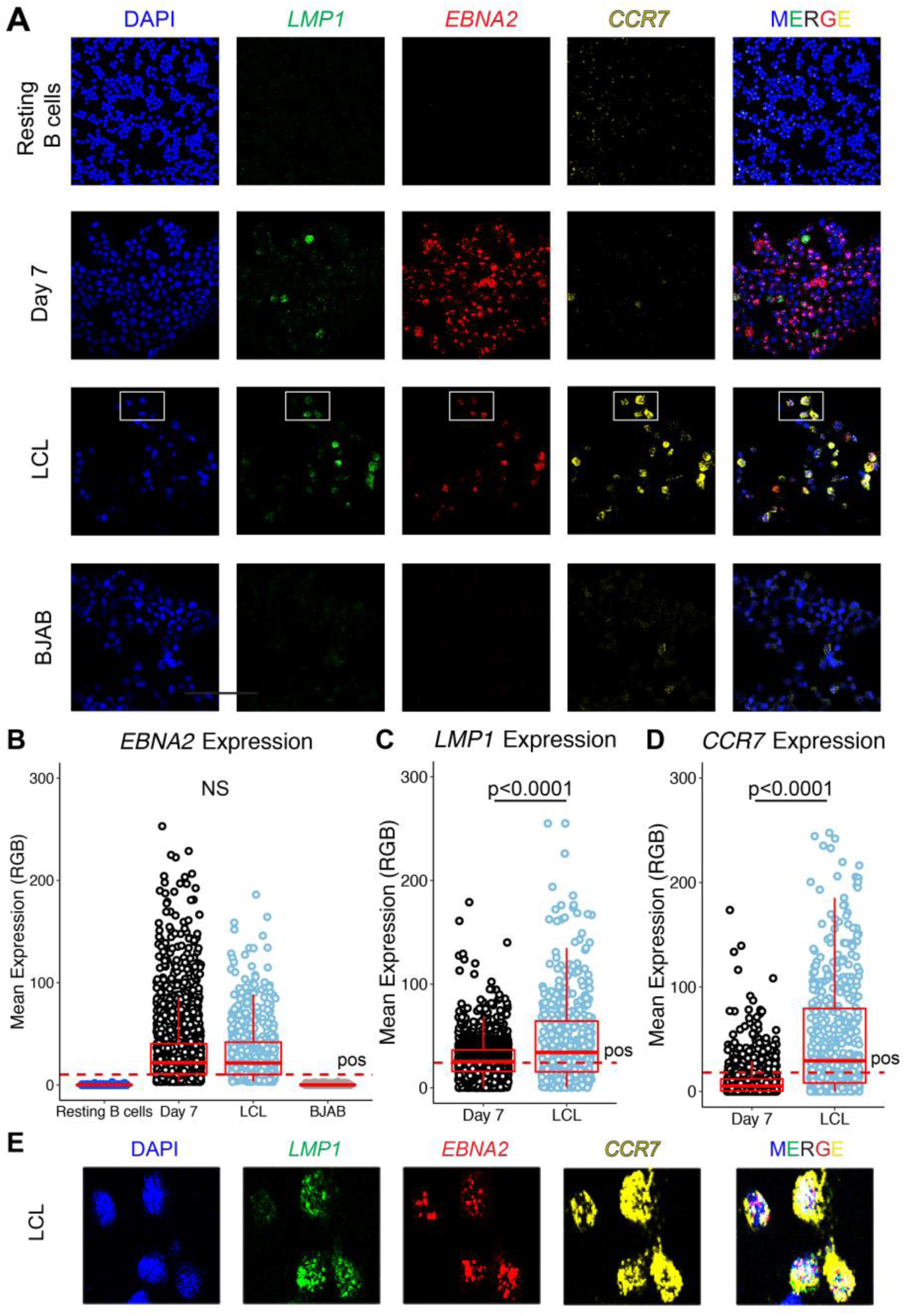
Multiplex RNA-FISH to distinguish EBV latency states. A) Confocal immunofluorescence images from Resting B cells, day 7 dpi EBV+ latency IIb proliferating B cells, LCLs and BJABs stained with LMP1, EBNA2 and CCR7 RNA FISH probes B) Quantification of EBNA2 expression. C) Quantification of LMP1 expression in EBNA2+ cells D) Quantification of CCR7 expression in EBNA2+ cells. E) Zoom in of boxed region in LCLs shown in A for EBNA2, LMP1 and CCR7 expression. For all quantifications, day 7 and LCLs are the average of 3 independent donors and seven fields of view. For resting B cells this is representative of one blood donor and 7 fields of view. BJAB also represents the average of 7 fields of view. * p<0.05, ** p<0.01, *** p<0.001 by one-tailed Mann-Whitney non-parametric t-test. All error bars denote SEM.

## DISCUSSION

In this study, we identified host biomarkers that distinguish EBV latency IIb from latency III. We recently established that the initial infection of primary human B cells with EBV displays a latency IIb phenotype where the viral EBNA proteins are expressed in the near absence of the LMPs (5, 20). However, later during primary infection latency III is observed where the EBNAs and LMPs are all expressed as in LCLs. Broad distribution of LMP1 within latency III populations makes these cells difficult to distinguish. As many EBV-positive tumors display cellular LMP1 heterogeneity (14, 15), it is important to determine whether these EBNA+/LMP1-cells are latency IIb or latency III as their immune recognition and response to chemotherapy may vary depending on viral gene expression (7, 10).

To address this question, we first confirmed that the NFκB induced surface protein, ICAM-1, is a proxy and reporter for LMP1-mediated NFκB signaling and LMP1 mRNA levels (19). We confirmed the broad range of LMP1/NFκB expression in latency III as observed by others (10, 13) and found a significant overlap of ICAM-1 surface levels between the latency IIb early-infected cells and ICAM-1^lo^ LCLs. Through FACS sorting coupled with RNA-sequencing, we found that the major determinant between LMP1^lo^ and LMP1^hi^ expressing LCLs is indeed NFκB signaling with a small component of cell cycle regulation through E2Fs. Importantly, we found that latency IIb gene expression profiles clustered distinct from latency III irrespective of LMP1 level or donor from which they were generated. Thus, latency IIb is a *bona fide* latency state.

We identified and validated four latency IIb and latency III specific host mRNAs that were differentially regulated between the states. *CCR6, FcRL4, FcRL3* and *TGFBR2* were specific to latency IIb while *CCR7, MGST1, DST* and *TSC22D3* were specific to latency III. Interestingly, CCR7 was one of the first genes demonstrated to be induced by EBV, originally being called EBI-1 (21). However, surface expression of CCR6 and CCR7 did not fully distinguish latency IIb from latency III. For this reason, we turned to multiplexed RNA-FISH to simultaneously measure LMP1, EBNA2 and CCR7 mRNA levels in single cells. This approach coupled with a multinomial regression model allowed us to predict whether an infected cell was latency IIb or latency III.

While protein-based expression analysis by immuno-histochemistry (IHC) is the gold standard in pathology labs, multiplex RNA-FISH is a promising new approach to decipher cellular heterogeneity in tumors (22, 23). This approach is equally sensitive compared to IHC but lacks the limitations of antibody specificity and sensitivity (24, 25). Indeed, multiplex RNA-FISH overcomes the issue of protein epitope variation or antigen retrieval by tiling probes for the target gene across the entire length of the mRNA. In our studies, multiplexing probes with distinct fluorophores enabled the detection of both viral and host mRNAs at single cell resolution.

EBV-positive lymphomas in immune-suppressed patients have been characterized to display latency III gene expression (EBNA+/LMP+) (20). However, several early pathology studies of EBV-positive PTLD and HIV lymphomas as well as more recent mouse models describe a significant EBNA+/LMP-cell population (14, 15). In a cord blood mouse model of EBV infection, EBNA2+/LMP1-cells were observed at a high frequency while latency III cells were rarely detected. This was hypothesized to be due to the increased immunogenicity of latency III cells (26). Similarly, a recent study using a mouse model of EBV/KSHV co-infection, latency IIb cells were detected at a high frequency (16). The pathophysiological relevance of latency IIb therefore supports our study to define latency-distinguishing host markers.

Autologous and allogeneic T-cell therapies targeting MHC-restricted viral antigens are used in the treatment of EBV-associated PTLD (27-33). These products are typically highly enriched for CD8+ cytotoxic T cells and therefore, proper EBV antigen presentation through MHC class I within these tumors is likely important for a robust clinical response. As LMP1 expression in a latency III B cell cycles between sub-populations that are LMP1^hi^ and highly sensitive to CD8+ T-cell killing and those that are LMP1^lo^ and much less sensitive (10), it will be important to distinguish whether EBNA2+/LMP1-cells within PTLD tumors are fixed LMP1^lo^ latency IIb cells or are latency III cells cycling between low and high LMP1 states. PTLD tumors with persistent latency IIb infection may be more difficult to treat with T-cell therapies than latency III-predominant PTLD. This remains to be tested by correlating EBV latency type in PTLD tissue with response to T-cell therapy.

Recent clinical studies have led to the development of LMP-specific CTLs for the treatment of EBV latency IIa tumors (LMP+/EBNA-) (34-36). For EBV-associated PTLD with predominately latency III gene expression, LMP-specific CTLs would be expected to have clinical efficacy and, indeed, a clinical trial is underway (37). In light of our findings regarding latency IIb, it remains pertinent to consider screening these tumors for LMP1 expression and perhaps excluding tumors that display a latency IIb expression phenotype. Coupling the host biomarkers that we have identified together with the viral EBNA2 and LMP1 using multiplex RNA-FISH could provide significant predictive power in screening these tumors for efficacy using T-cell therapies and chemotherapeutics.

## MATERIALS AND METHODS

### Cell lines, culture conditions, and viruses

Peripheral Blood Mononuclear Cells (PBMCs) were obtained from whole blood from the Gulf Coast Regional Blood Center (Houston, TX) via centrifugation over a Ficoll Histopaque-1077 gradient (Sigma, H8889). The B95-8 strain of Epstein-Barr Virus was generated from the B95-8 Z-HT cell line as previously described (38). Virus infections were performed by adding either 100 μL of filtered B95-8 Z-HT supernatant to 10^6^ PBMCs or 500 μL of B95-8 Z-HT per 10^6^ B-cells, as determined by FACS.

Cell lines were cultured in RPMI 1640 media supplemented with 10% (LCLs) or 15% (primary B cells) fetal bovine serum (FBS) (Corning), 2 mM L-Glutamine (Invitrogen), 100 U/mL penicillin, 100 μg/mL streptomycin (Invitrogen). All cells were maintained at 37°C in a humidified incubator with 5% CO_2_.

### Flow Cytometry and Sorting

To track cellular division, cells were stained with CellTrace Violet (Invitrogen, C34557), a fluorescent proliferation-tracking dye. For analytical panels, 10^6^ PBMCs on day 7 post-infection with EBV B95-8 or 10^6^ LCLs were washed once with FACS buffer (PBS + 5% FBS) and stained with the following antibodies in isolation or in combination for 30-60 minutes in the dark at 4°C: ICAM-1 PE (BioLegend, #353106), CCR6 PE/Dazzle (BioLegend, 353430), CCR7 PE/Dazzle (BioLegend, 353236), or CD19 APC (BioLegend, 302212). After incubation, cells were washed once with FACS buffer and 10,000 blank counting beads (Spherotech, #ACBP-50-10) were added to each tube. Data was collected on a BD LSRFortessa and 1000 blank beads were used as the stopping gate. All samples were stained and analyzed via FACS at the same time to ensure consistency in analysis. Marker positivity was determined using matched fluorescence minus one control.

For sorting experiments, proliferating cells were sorted to purity using CD19 APC (BioLegend, 302212) positivity as well as a dilution of CellTrace Violet (CD19^+^/CTV^lo^) on a MoFlo Astrios Cell Sorter at the Duke Cancer Institute Flow Cytometry Shared Resource. LCLs were sorted to purity after staining with ICAM-1 PE (BioLegend, #353106) and gated for the bottom, middle and upper 15% of ICAM-1 expressing cells.

### RNA-Sequencing and Analysis

Whole RNA from sorted early EBV infected latency IIb B cells and from sorted donor matched LCLs were isolated using a RNeasy-Kit (Qiagen, #74104). mRNA sequencing libraries were prepared using a Kappa Stranded RNA-Seq Library Preparation Kit (Kappa Biosystems, KR0934) and sequenced on an Illumina Hiseq 4000 at the Duke University Sequencing and Genomics Shared Core Facility. Resulting single-end, unpaired reads were aligned to the human genome (hg38) using Hisat2 (39). Resulting BAM files were converted to SAM files using samtools and transcripts were assembled using Stringtie. Assembled transcripts were quantified using the R package Ballgown. Normalized RPKMs were exported from Ballgown and used for heatmap visualization and log_2_(RPKM+1) calculations. Statistical significance and false-positivity were determined using ballgown. Heatmaps were generated using Morpheus from the Broad Institute (https://software.boradinstitute.org/morepheus) and similarity matrices were created using the R package pheatmap (https://CRAN.R-project.org/package=pheatmap). RNA-sequencing coverage maps were generated using UCSC Genome Browser in a Box (GBiB) (40).

### RNA Isolation, RT-qPCR and Primers Used

Total RNA from sorted EBV infected early latency IIb proliferating B cells or sorted LCLs was isolated using an RNAeasy kit (Qiagen, #74104) according to the manufacturer’s instructions. One microgram of total RNA was reverse transcribed into cDNA using the High-Capacity cDNA Reverse Transcription Kit (Applied Biosystems, 4368814) according to the manufacturer’s instructions. Resulting cDNA was diluted in ultra-pure H_2_O and 5 ng per reaction was used for RT-qPCR via SYBR green (Quantabio, cat# 95054) detection method. Relative expression was calculated using the ΔΔCT method using SETDB1 as an endogenous control. The following table lists all primers used for RT-qPCR in this study:

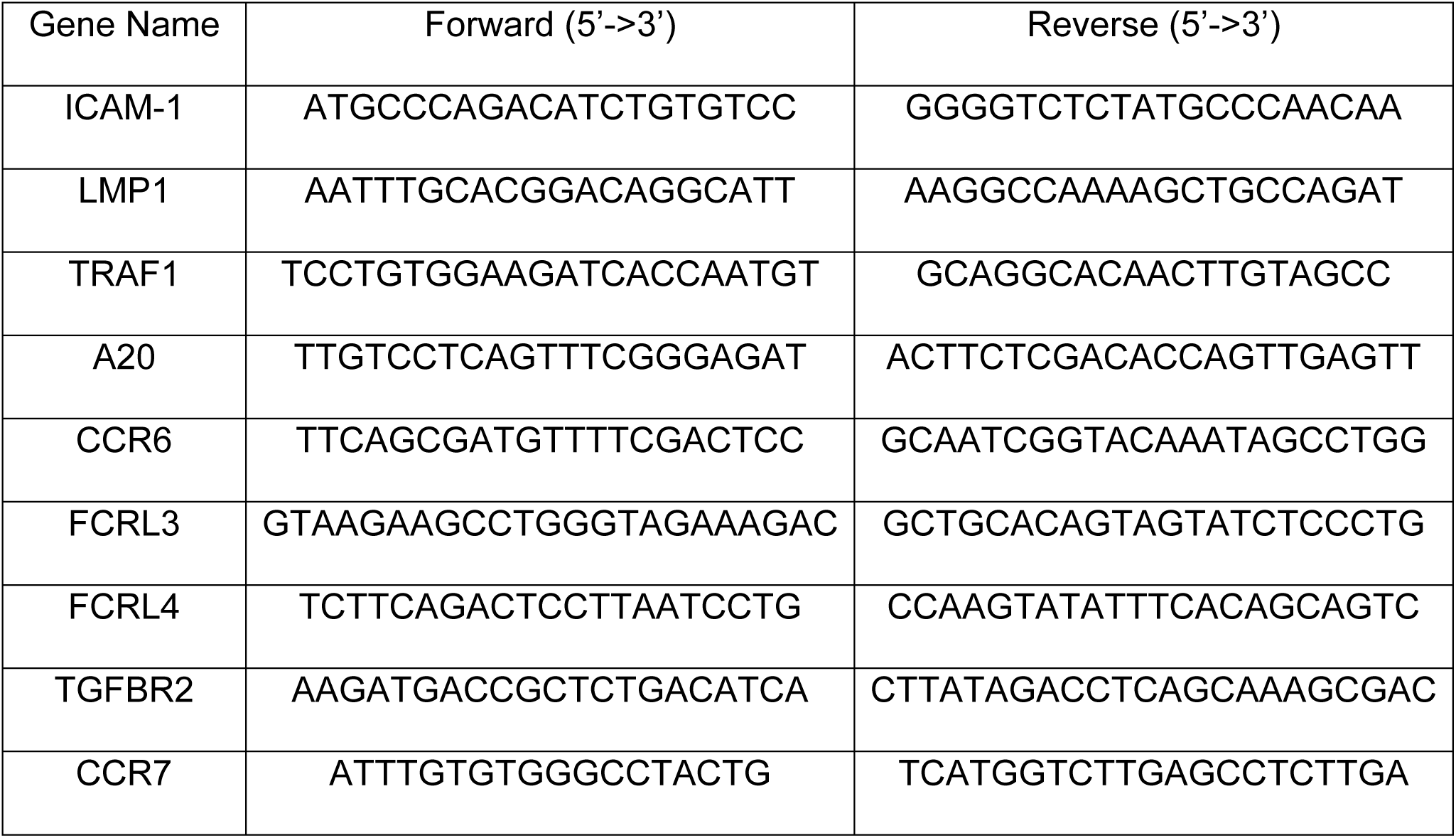
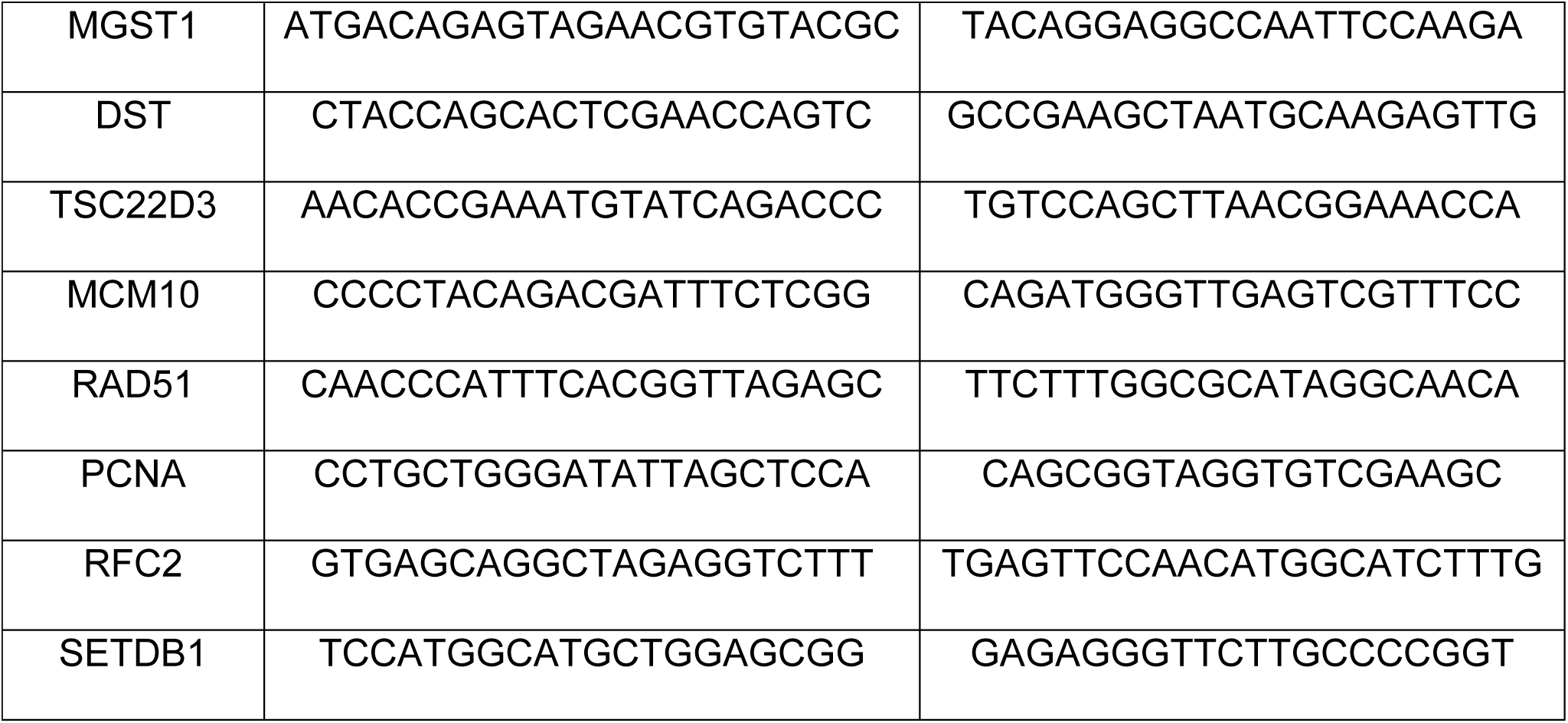

### RNA-FISH

RNA-FISH was conducted using the Advanced Cell Diagnostics (ACD) RNA SCOPE Multiplex Fluorescent v2 Assay (Advanced Cell Diagnostics, #323100). In brief, resting B cells isolated from peripheral blood (BD IMAG Human B Lymphocyte Enrichment Set – DM, BD #558007), sorted latency IIb proliferating B cells on day 7 dpi and LCLs were washed once in PBS, fixed in 10% neutral buffered formalin for 1 hour at 37°C, washed again in PBS and resuspended in 70% EtOH before being cytospun onto glass slides using a Cyto-Tek Sakura table-top cytofuge at ∼735*g* for 22 mins. Slides were dried for 20 mins before being fixed in an ethanol gradient of 50%, 70% and 100% EtOH for 5 mins each. Slides were stored overnight at −20°C in 100% EtOH before being dried and having a hydrophobic barrier applied to the slide using an ImmuEdge pen (Vector, H-4000). Samples were first treated with peroxide for 10 mins at RT to quench endogenous peroxidase. After peroxide treatment, cells were treated with ACD protease III for 30 mins at 40°C before proceeding to the standard RNA-SCOPE multiplex fluorescent V2 assay protocol, performed to the manufacturer’s instructions. Cells were stained with a probe mixture containing HHV4-LMP1-C1 (Advanced Cell Diagnostics, ref#414681), HHV4-EBNA2-C2 (Advanced Cell Diagnostics, ref#547771-C2), and either Hs-CCR7-C3 (Advanced Cell Diagnostics, ref#410721-C3). After hybridization the signal was amplified and conjugated to either Fluorescein, Cy3, or Cy5 Perkin Elmer TSA Secondaries (Perkin Elmer NEL741E001KT, NEL744001KT and NEL745001KT, respectively). Slides were stained with DAPI before being mounted with ProLong Gold antifade (Invitrogen, P10144). Slides were dried for 30 mins at RT before being moved to 4°C for long-term storage. All images were acquired on a Zeiss 780 Upright Confocal Microscope and resulting images were analyzed with Fiji.

### Fiji Image Analysis

Images were processed using in-house Fiji macros. The macro performs the following functions: For each sample the DAPI image and corresponding fluorescent channel image are simultaneously imported into Fiji. A Gaussian Blur (σ = 2) is applied to the DAPI image and then an Otsu threshold is applied. The DAPI image is then converted to binary and the watershed function is then applied to distinguish potentially overlapping nuclei. A threshold is then applied to the fluorescent channel image (auto for fluorescein and Cy5, minimum for Cy3). The DAPI image is subsequently selected and the Fiji Set Measurements window is utilized to report the area, mean, minimum and maximum intensity redirected to the fluorescent channel image. Fiji Analyze particles is then used to determine the intensity of foci in the fluorescent channel image that lie within boundaries identified by the DAPI channel image.

Once the macro had been applied to all images for all fluorescent channels all of the raw data is curated. The expression of day 7 cells and LCLs stained with positive and negative control probes provided by the manufacturer (Advanced Cell Diagnostics, REF#321801 and 321831, respectively) were plotted on a histogram and positivity thresholds were set at the point where the positive and negative control histograms intersect (data not shown). For EBNA2-Cy3, a minimum threshold is used allowing for harsh discrimination between EBV+ and EBV-cells. For LMP1-fluorescein and CCR7-Cy5, to be more tolerant of “low” expression, a less harsh “automatic” thresholding method was used.

Having established cutoffs for EBNA2, the data was subsequently curated to only report LMP1 and CCR7 expression in cells that were positive EBNA2 mRNA to ensure that we analyzed only EBV infected cells. Due to the non-gaussian distribution of this positive signal, a Mann-Whitney non-parametric t-test was used to determine statistical significance.

## DATA AVAILABILITY

The RNA-sequencing data will be uploaded to Gene Expression Omnibus (GEO) under accession number XXXX.

## ACKNOWLEDGMENTS

This work was supported by National Institute of Health (NIH) grants R01-DE025994 (to M.A.L.), T32-CA009111 (to J.E.M., J.D and A.M.P.), and F31-CA180451 (to A.M.P.) and F31-DE027875 (to J.D.). Additional funding came from the Duke CFAR, and NIH-funded program (grant 5P30-Al064518).

The authors would like to thank Dr. Michael Cook, Nancy Martin, Lynn Martinek and the Duke Cancer Institute Shared Flow Cytometry core facility for invaluable assistance with flow cytometry and cell sorting. We would also like to thank Dr. Lisa Cameron, Dr. Yasheng Gao and the Duke Light Microscopy core facility for help with confocal microscopy. We would also like to thank the Duke Sequencing and Genomic Technologies core facility for help with mRNA sequencing. Lastly, we would like to thank Dr. Dirk Dittmer, Justin Landis, Dr. Lorenzo Zaffiri, Dr. Elizabeth Pavlisko, Francine Kelley, Andrew Nagler, Dr. Scott Palmer and Daniel Leibel for helpful discussions during the drafting of this manuscript.

## AUTHOR CONTRIBUTIONS

J.E.M, A.M.P., and M.A.L. conceived and designed research experiments. J.E.M. performed buffy coat isolations from peripheral blood, EBV infections, all bioinformatic analysis of RNA-sequencing data, RT-qPCR validation of RNA-sequencing, FACS analysis of RNA-seq targets, and RNA-FISH. J.D. and L.J.S. prepared mRNA sequencing libraries. A.M.P. performed the initial infection, sorting, and RNA isolation. J.E.M. and M.A.L. wrote and all authors edited the manuscript.

## Supplementary Data Legends

**Supplemental Table S1: List of genes downregulated from latency IIb to ICAM-1**^**lo**^ **LCL**

**Supplemental Table S2: List of genes upregulated from latency IIb to ICAM-1**^**lo**^ **LCL**

**Supplemental Table S3: List of latency IIb and latency III specific genes**

